# An Open-Source Image Analysis Method for Quantifying Reporter Fluorescence

**DOI:** 10.1101/2025.02.02.636071

**Authors:** Kathryn F. Ball, Jacob A. Perez, Allen M. Cooper, Charmaine U. Pira, Kerby C. Oberg, Christopher G. Wilson

**Author notes:** **Correspondence:** Kathryn F. Ball.

## Abstract

Image analysis is a rapidly developing field that provides unique opportunities to characterize and quantify spatial information in images. We study *cis*-regulatory modules (CRMs), non-coding DNA regions that regulate gene expression, during development using fluorescent reporters *in vivo.* Characterizing CRM activity during development presents challenges including image segmentation into biologically relevant regions of interest that are not easily distinguishable via common segmentation methods, fluorophore band passing, and variable transfection undermine standardized analysis. To quantify and compare CRM activity levels, we compiled an open-source computer vision tool stack in the form of a *Python*-based *Jupyter* notebook and tested four analysis methods to assess their efficacy in quantifying CRM expression in limb development. This *Jupyter* notebook provides a reproducible, standardized workflow that can be adapted to numerous image analysis applications.

## 1 Introduction

### 1.1 Background

Fluorescent markers have long been used to report the activity, location, and spatial relationship of molecules in cells and organisms. This can be done through a variety of techniques such as protein tagging, fluorescent immunohistochemistry, or fluor-labeled antisense probes to determine cell location, colocalization of components, or even inter- and intra-molecular interaction (1,2). Fluorescence probes allow us to detect structures that might otherwise be very difficult to detect or invisible with light microscopy. As imaging technologies and their methodological applications have advanced, microscopy has moved from qualitative and descriptive to more quantitative and determinant. The field of image analysis has paralleled this path and adopted quantitative methods. Current image analysis practices include a host of computational tools and digital imaging methods that allow for relatively easy image quantification, including bright field, fluorescent, MRI, CT, etc. However, most image analysis protocols lack standardization—this is understandable due to the wide variety of imaging applications in the biological sciences.

Recent advances in genomic analysis have facilitated investigations into *cis*-regulatory modules (CRMs), regulatory sequences that control the temporal and spatial expression of genes. To evaluate and annotate the functional domains of CRMs, fluorescent reporters are used to detect subtle changes in activity with variations in sequences. Our goal in this paper is to provide a workflow that can extract quantitative data from fluorescence images despite variations in transfection efficiency and bleed-through from other fluorophores and provide a step-by-step protocol to maximize reproducibility and transparency.

### 1.2 Review of Available Software Methods and Limitations

Proprietary image analysis tools such as *Imaris Bitplane* or *Visiopharm* have been used for many years and each have their own underlying assumptions (platonic solid problems versus corpuscle under-estimation) that may bias measurements. Open-source tools provide better transparency and reproducibility than their commercial counterparts, typically with more rapid bug-fixes and security patches than the proprietary tools. Perhaps the most notable open-source tool for image analysis is *ImageJ* (Schneider 2012). When Wayne Rasband developed *ImageJ*,’s predecessor *NIH Image* in 1987 it revolutionized the world of image analysis and has become perhaps the most used image analysis tool in science. *ImageJ* was ubiquitous and freely available, meaning that, for the first time, individual investigators could create their own custom tool stacks for image analysis—akin to the tool stacks that are used for omics analysis.

Despite its benefits, *ImageJ* has some limitations. The modern version of *ImageJ/ImageJ2/FIJI* is written in *Java*, which has a syntax that is deliberately like *C/C^++^*. This is an advantage from a computer science perspective since it makes *ImageJ* functional across multiple operating systems and hardware platforms but can be considered a disadvantage from the perspective of a biologist who is new to programming. *Java,* like *C^++^*, is not user-friendly for those who come to the language without a strong programming background. *Java* was developed for portability but has a reputation for being slower than *C/C^++^* and *Java* executables take up more memory. Further, *ImageJ2* limits image size to 2 gigapixels. We note that, for those who would like to use *Python* scripts for *ImageJ*, the *PyimageJ* wrapper (3) may be useful.

*Python* is a widely used general programming language, known for its relatively simple syntax, user-friendly interface (in the form of the *iPython* shell or *Jupyter* notebook), making it a useful programming language and tool for scientists, not already well-versed in computer languages. In 2018, a survey of 206 scientists and researchers showed that *Python* and *R* were the most-used languages (used by 67.0% and 59.2%, respectively), while *C^++^* trailed at 26.2% and *Java* was only used by 17.0% of those surveyed (4). With the addition of high-performance matrix math (*NumPy*) and scientific function (*SciPy*) libraries, *Python* performs at speeds comparable or faster than *MATLAB*, one of the most common engineering and scientific computing languages (5), when using optimized math libraries. *Python* executables also occupy more memory but, in our opinion, the compromise for ease of coding and clear syntax make it a better choice for the average biologist learning to address a specific analysis problem and deal with their data.

*Python* offers ease of use and allows users to process images with computer vision modules such as *PIL/Pillow OpenCV*, and *scikit-image* and works with modules such as *TensorFlow* and *Keras* for machine learning applications. Several open-source image analysis software packages are available for use (Table 1), but we found none that met our needs in terms of image segmentation and are designed to work with a standard two-fluorophore fluorescent experiment.

**Table 1.**

| Name | Language(s) | Key Functions | Citation | Link |
| --- | --- | --- | --- | --- |
| SamuROI | Python | Multiple ROI analysis at different scales, TIF-based timeseries | (16) | https://github.com/samuroi/SamuROI |
| SIMA | Python | Time-series fluorescence analysis | (17) | https://github.com/losonczylab/sima |
| ACQ4 | Python | Acquisition from different hardware types and analysis | (18) | http://www.acq4.org/ |
| μManager | C++ (has Python wrapper and can interface with MATLAB) | Microscope hardware interface, acquisition, and analysis | (19) | https://micro-manager.org/ |
| PathML | Python | Whole-image slide deep learning image analysis | (20) | https://github.com/markowetzlab/pathml |
| Orbit | Python | Whole slide image analysis with ML | (21) | https://www.orbit.bio/ |
| ImageJ | Java | General-purpose image analysis | (22) | https://imagej.net/ij/index.html |
| Qupath | Java, C | Pathology and whole slide image analysis | (23) | https://github.com/qupath/qupath |
| Icy | Java | Bioimage informatics platform | (24) | https://github.com/lcy-imaging |
| Ilastik | Python, C++, Shell, Java, Jupyter Notebook | ML for cell segmentation, classification, and tracking | (25) | https://github.com/ilastik/ilastik |
| Cell Profiler | Python, Java, Makefile, Jupyter Notebook, Shell | Cell and bioimage analysis pipelines | (26,27) | https://cellprofiler.org/<br>https://github.com/CellProfiler |
| Cytomine | Python, Java, Javascript | multi-gigapixel imaging data | (28) | https://github.com/cytomine<br>https://doc.cytomine.org/ |

### 1.3 Review of Available Laboratory Methods and Limitations

Many methods that complement qualitative data such as flow cytometry or RT-qPCR rely on disassociating cells or cellular structures and thus lose important spatial information in the process. New methods have recently become available for multiplexing gene expression while maintaining distinct spatial information such as HCR RNA-FISH. While this method provides exciting new opportunities for gene expression, it cannot report on CRM activity since it is based on antisense probes to mRNA and, like all *in situ* hybridization, is limited in its quantitative application by variation in probe specificity, melting temperature, and the length of staining and wash steps.

### 1.4 Methodological Proposal

Our aim was to develop a protocol for quantitative comparisons across datasets to address the following gaps in available image analysis workflows: high-throughput batch processing, account for band-passing, limit user bias, and collect fluorescence intensity without losing information about relative position and orientation. To fill these gaps, we developed a *Jupyter* notebook that allows us to standardize image acquisition and processing as much as possible within the context of our laboratory’s work. We propose an image analysis workflow including standard steps such as grayscale conversion, noise removal, and image segmentation (see (6) for review), and additional steps for data cleaning and organization, bleed-through removal, and dynamic ROI selection with code blocks for optimizing each of the steps. In an effort to make this open-source tool stack available to others, we present our step-by-step protocol with explanatory text, in this paper and the associated *Jupyter* notebook which is available at https://github.com/KateBall/Quantitative_Image_Analysis under the GNU Public License (GPL, ver. 3).

## 2 Materials and Equipment

### 2.1 Targeted Regional Electroporation (TREP)

Chicken embryos were staged according to the Hamburger and Hamilton (HH) method (14). The embryonic coelom of the lateral plate mesoderm of stage HH14 embryos was injected with DNA solution (2 µg/µL pTK-ZRS-EGFP, 0.2 µg/µL pCAGGS-RFP with Fastgreen and Tris-EDTA buffer). Plasmids were electroporated into the presumptive limb using the CUY-21 Electroporator (Protech International Inc., Boerne, TX) as previously described (15). Embryos were incubated for 48 hours post-electroporation then harvested. We visualized fluorescence with a Leica MZ FLIII fluorescent stereo microscope using 41012 HQ:FLP FITC/EGFP and 10446365 TXR filters (Chroma Technology Corp., Brattleboro, Vt.); images were captured with a Sony DKC-5000 camera and acquired using *Adobe Photoshop* (version 6.0).

## 3 Methods

### 3.1 Case Study (ZRS)

We study CRMs active during limb development. As limbs develop, they form a bud that emerges from the body wall (Fig. 1A). The zone of polarizing activity (ZPA) is a small cluster of mesenchymal cells in the posterior aspect of the limb bud that express Sonic Hedgehog (Shh) (Fig. 1A). Shh directs development along the anterior-posterior axis of the limb and is responsible for digit patterning. The ZPA Regulatory Sequence (ZRS) is a CRM found in non-coding DNA that is conserved across vertebrates and regulates Shh expression in the limb. To monitor CRM activity, we electroporate the presumptive limb region of chicken embryos with a mixture of control and experimental plasmids (Fig. 1B, see appendix for technical details). Control plasmids contain red fluorescent protein (RFP) driven by a cytomegalovirus (CMV) enhancer linked to a β-actin promoter, while experimental plasmids contain our enhancer of interest and a green fluorescent protein (GFP) reporter driven by a minimal thymidine kinase (TK) promoter. RFP is constitutively expressed and serves as a positive transfection control, while GFP will be expressed only in tissue with the transcription factors necessary for enhancer activity. Quantitative image analysis for these experiments presented certain challenges, namely RFP bleed-through into GFP images due to spectral overlap (Fig. 1C-D), transfection variability due to the nature of electroporation (Fig. 1D), and the need for a standardized method of normalization. Thus, we present our analysis method, a flow-chart highlighting the key steps is shown in Fig. 2.

**Figure 1.**
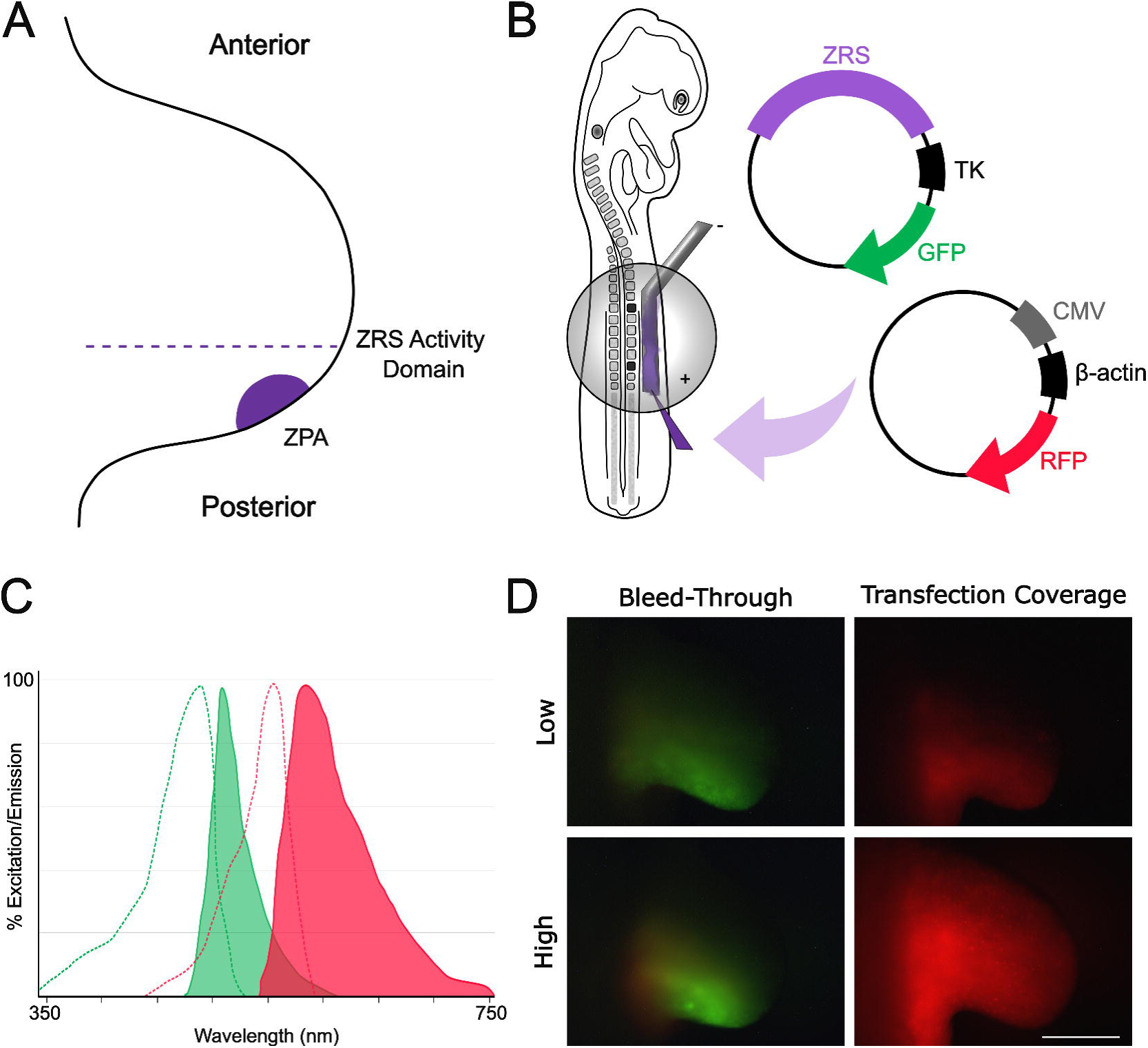
ZRS is a limb-specific enhancer/Bleed-through and transfection variation present challenges to image analysis. A) Dorsal view of a developing limb bud, Posterior (below dotted line), and ZPA region (shaded purple). B) Hamburger-Hamilton stage (HH14) chicken embryo receiving plasmids via electroporation. C) Excitation (dotted lines) and emission (shaded peaks) spectra of GFP (green) and RFP (red). Bleed-through results from overlap of fluorochrome excitation and emission spectra. D) Left: GFP images with relatively low (top) and high (bottom) levels of bleed-through. Right: Limbs electroporated with CMV-β-actin-RFP demonstrate transfection variability. Scale bar: 500μm. Images have been cropped to remove empty space.

**Figure 2:**
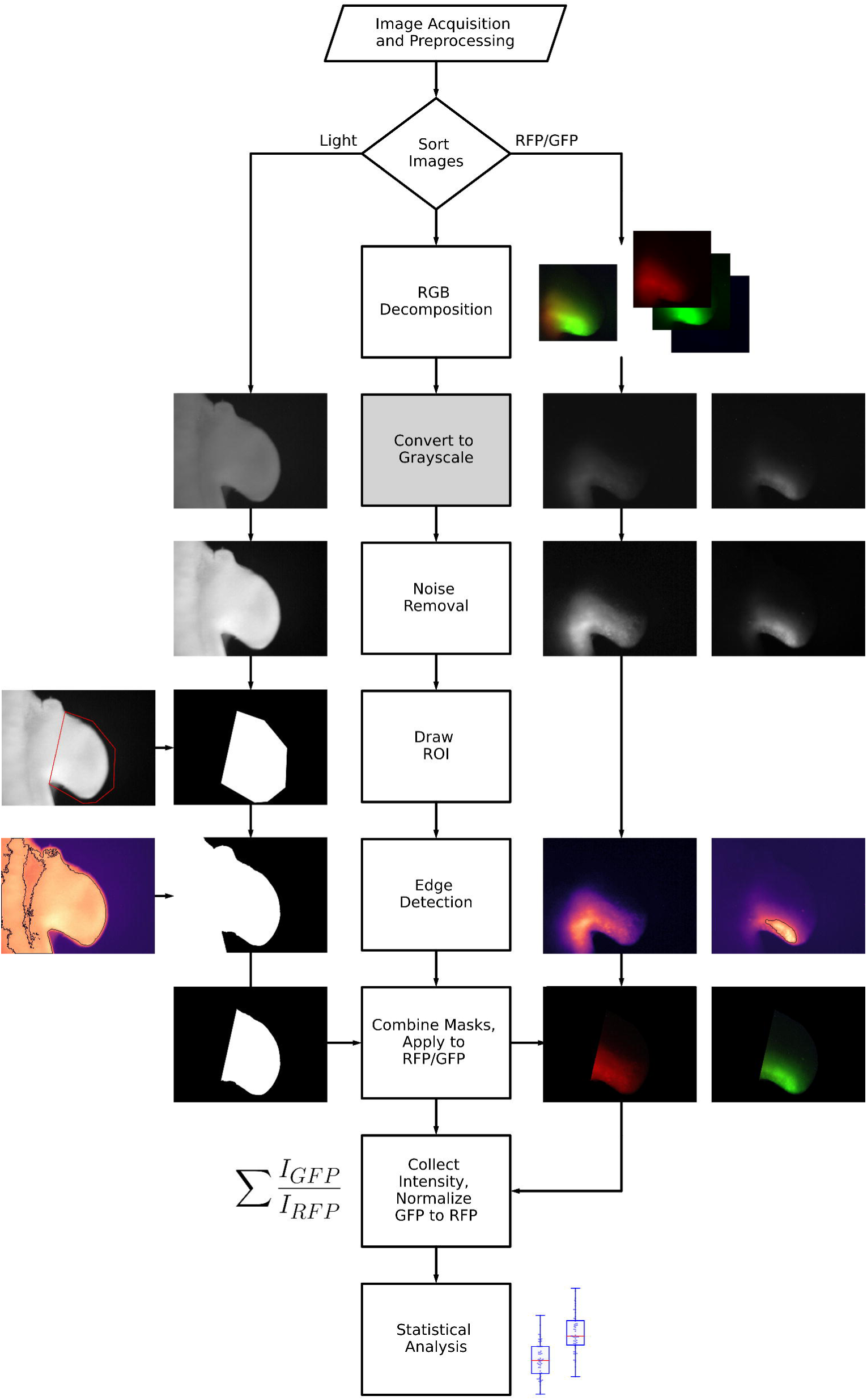
Method Flowchart. Graphical overview showing the computational steps in the given workflow. Image acquisition and preprocessing includes TREP and protocol steps 0-3. GFP image with a high amount of bleed-through is shown for the RGB decomposition example, while the remaining steps are demonstrated with example settings performed using the light, RFP, and GFP images for a single limb. Fluorescent images (RGB decomposition and masked RFP and GFP) in this figure have been adjusted for lightness (30%) and contrast (10%) using Inkscape 1.2.2. Flow chart made with Schemdraw© 0.17.

#### 3.1.1 Protocol

##### Step 0: Load Dependencies

Loads the *Python* modules needed for the notebook. See the *Jupyter* notebook for more information about each module used, and links to library resources. Some of the modules loaded are only used in the initial testing stages and not in the final workflow.

##### Step 1: File Renaming and Organization

Since the software used for image acquisition does not automatically store metadata such as exposure time, magnification, or fluorescent filter used, we stored this information in the filename. Since our file nomenclature evolved over time, we retroactively renamed files to fit the regex format: `‘^(\d{4})-(\d{2})-(\d{2})_E(\d{2})_(.+)_(.+)_RLD_63x_([a-zA- Z]+)_([0-9]+)s\.(jpg|JPG|png|PNG|tif|TIF)$’`. This format distinguishes between files’ experimental date `‘^(\d{4})-(\d{2})-(\d{2})_’`, embryo number within the experiment `‘E(\d{2})_’`, enhancer species and construct code `‘(.+)_(.+)_’`, limb and view `‘RLD_’` (right limb dorsal view), magnification `‘63x_’` (total magnification), filter used `‘([a- zA-Z]+)_’`, exposure time `‘([0-9]+)s’`, and file format `‘\.(jpg|JPG|png|PNG|tif|TIF)$’`. Other naming formats could be used, so long as they are uniform and distinguish between key experimental parameters. Although this filename format is lengthy, it reduces the number of directories needed to keep the files organized.

In this workflow, we acquired images using an older image acquisition system so information surrounding the image, such as date taken, embryo ID, and exposure time, is stored in the filename and not the metadata. Ideally, users of our workflow will have access to an image acquisition system that stores metadata which can be archived as embedded information rather than requiring separate notation or user modification of the filename. All filenames must be converted to a uniform nomenclature to use with this notebook. The authors strongly suggest that lab groups establish a common file naming and directory structure convention. We recommend all files contain a date following the ISO-8601 date/time format and nomenclature that adheres to ISO-9660 standard and avoid the use of spaces in filenames and directories.

##### Step 2: Select Files and Make Figures

In this example, for each embryo there were three pictures, a brightfield (‘Light’) image, one taken with an RFP fluorescent filter (Texas red), and one taken with a GFP fluorescent filter (41012 HQ-FLP). While images with multiple exposure times were taken with each filter (0.5s, 1s, 2 for Texas Red, and 2s and 4s for 41012 HQ-FLP), only one exposure time for each is used in the final process. We selected 0.5s for the RFP image since transfection can vary and many of the 1s images are oversaturated. For the GFP image we used the 2s exposure time since the 4s image often has a high background. We used the Regex mentioned above to select embryos of the desired construct and RFP and GFP exposure times. We excluded embryos that did not meet our inclusion criteria (missing the appropriate exposure time: ‘_excld’, poor morphology: ‘morphexcld’, poor transfection (RFP): ‘redexcld’, or contains high levels of autofluorescence: ‘autofluor’) using an ‘if’ statement to recognize these keywords in the directory. Once an image meets the selected format and criteria, we add its file path to a list for further processing.

To visually inspect the embryos and ensure that for each embryo there was exactly one Light, one RFP and one GFP image, we created an iterative function “make_figure” that would display the limb images along with their experimental date and embryo number.

##### Step 3: Histograms

To identify oversaturation and determine the distribution of pixels, which will be used later in setting thresholds, we inspect each image’s pixel data either collectively using the histogram_basic function, or of each RGB channel separately with the histogram_rgb function. Limbs with fluorescent images that are oversaturated are excluded from further analysis.

##### Step 4: Bleed-through Removal

Due to the overlap of excitation and emission spectra, the signal of different fluorophores can be detected despite the use of specific fluorescence filters (Fig. 1D). We found that our positive transfection control, RFP driven by a CMV enhancer-β-actin promoter, bleeds through the green filter and is visible in the GFP image. Since the goal of this method is to measure experimental reporter activity, we needed to separate the control light from the experimental light. We considered different color-spaces as a way of identifying and handling the bleed-through. While HSV is commonly used for object identification, it did not perform well at segregating the bleed-through. Ultimately, we decided to decompose the RGB channels to isolate the source of the bleed-through. We created a null array of the same shape as the RGB image, sliced the red, green, and blue (RGB) values (Fig. 3A) and made three new images, each with the values for one of the RGB channels and the other channels filled with the null array (Fig. 3B). We then measured the intensities for all the pixels in each decomposed image (Fig. 3C). Band-passing in fluorescence images can be avoided by carefully selecting fluorophores that have minimal excitation/emission overlap and using high-quality fluorescent filters that are specific to your fluorophores. While our group is making that change for the future, we used this computational method to process pictures already acquired.

**Figure 3:**
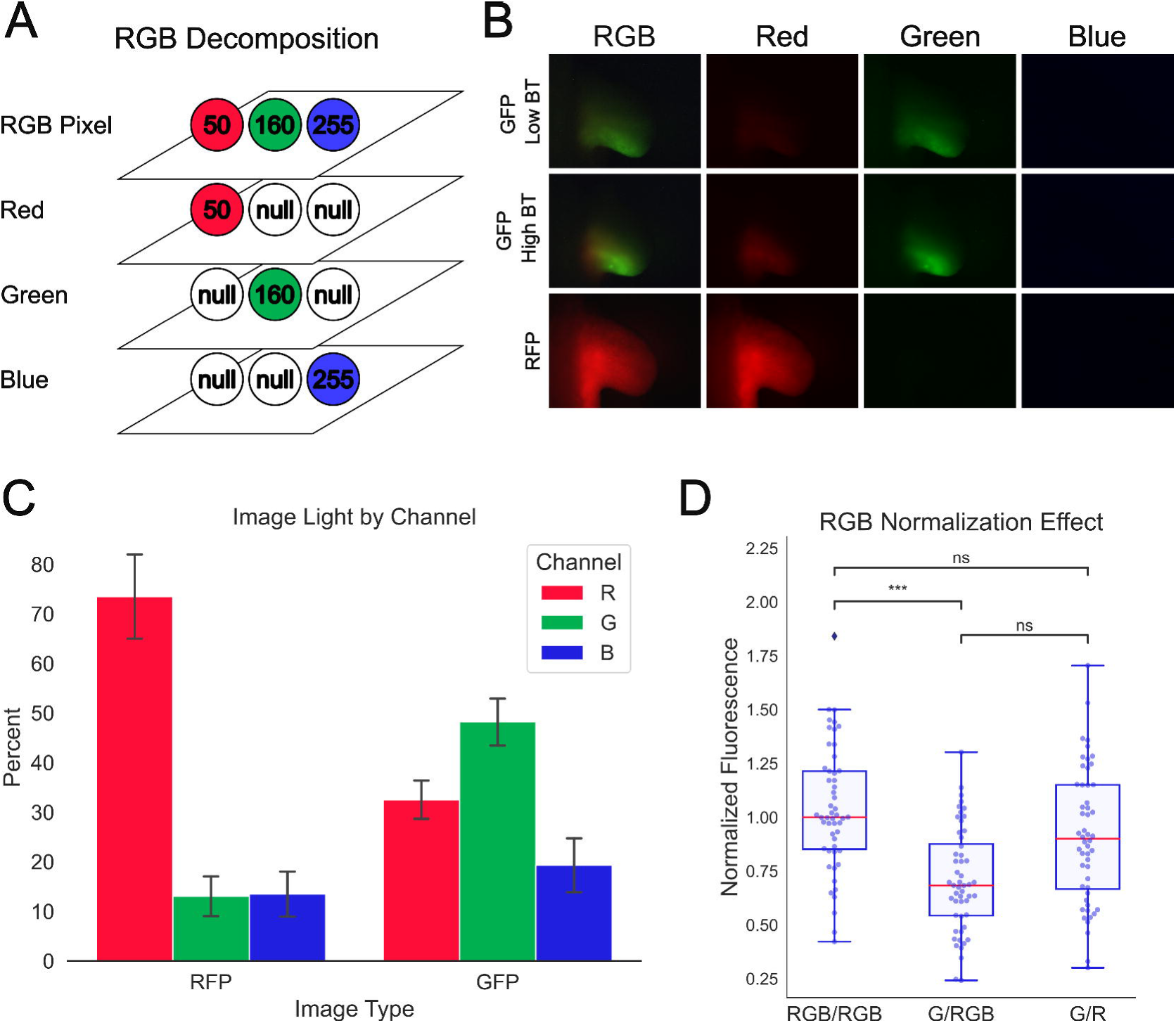
GFP images contain a proportionally high amount of light in the red channel. A) Diagram of the RGB values in a single pixel, and the corresponding pixels in each of the resultant single-channel images. B) Example decomposed images: GFP images with relatively low (top) and high (middle) levels of bleed-through (BT). RFP (bottom) shown for comparison. C) Bar chart representing the percent intensity in each color channel out of total intensity for all the pixels in each decomposed image (error bars: s.d.). D) Boxplots representing the three methods for collecting and normalizing the fluorescence data (Tukey’s HSD, ***: p<0.001, ns: not significant, n=52).

##### Step 5: Convert to Grayscale

This step combines the values from the red, green, and blue color channels into a single monochrome value on a 0 to 1 scale. Having all intensity information on a single scale is necessary for downstream steps such as noise reduction and thresholding. Different conversion tools assign different weights to each of the color channels. We tested both *OpenCV* and *scikit-image* libraries and for the sake of consistency chose to use commands from within the same module to read image files, convert to grayscale, and filter.

##### Step 6: Noise Removal

Image noise can arise from artifacts in the sample or objective lens, dead camera pixels, or other sources (7). To reduce the noise, the image can be passed through a low-pass filter technique called blurring. We compared the *OpenCV* methods: Average, Median, Gaussian, and Bilateral blurring filters. For light images, most blurring filters behaved similarly, but for fluorescence images the Average and Gaussian blurs (each with a 5×5 kernel) failed to remove noise and the Median blur removed noise, but decreased the contrast between signal and background, which could affect edge detection. The Bilateral filter is a kind of Gaussian blur designed to preserve edges and it performed the best, consistently removing noise and maintaining the difference between signal and background. For the *scikit-image* library we tested the Median, La Place, Gaussian, and Bilateral blurring filters. The La Placian filter did not work for any images. The Median, Gaussian, and Bilateral filters all performed well, though the Bilateral filter seemed to do a slightly better job of removing noise and maintaining contrast. Overall, we decided to use the *OpenCV* Bilateral filter, as it worked well, though the *scikit-image* Bilateral performed just as well.

To account for non-specific fluorescent labeling, background noise, and motion artifacts, some researchers perform additional background subtraction (8–10). We created a mask of ⅓ the image height by ⅓ the image width to collect background fluorescence in each image and compared normalized intensity values with and without background subtraction and found that background subtraction did not significantly affect the outcome.

##### Step 7: Edge Detection

Contours and thresholding are powerful tools to detect the edges of objects within an image. We compared contour methods from the *Matplotlib*, *OpenCV*, and *scikit-image* modules. The *Matplotlib* contour function is good for visualizing various thresholds within an image and highlights what regions have a similar intensity level.

We compared numerous thresholding methods available in *OpenCV* and *scikit-image*. Most thresholding methods consistently detected the edge of limbs under regular light (Global and Adaptive Mean from *OpenCV* and Isodata, Li, Mean, Minimum and Triangle from *scikit-image*) though the Li, Minimum, and Triangle methods pick up more noise and Yen often does not work at all. We chose to use the Otsu method because it does the job while avoiding noise. The Isodata and Mean methods may be good alternatives. For fluorescent images that are bright, the Minimum method does not pick up anything and the Yen method only works half of the time. The Mean method usually has a high background, the Isodata, Li, and Otsu (11,12) methods seem to do a pretty good job, and the Otsu method appears to have the lowest background. The Triangle method performs similarly to the Otsu method, but in some cases, it is too stringent and does not pick up a large area of weak fluorescence if there are a few brighter foci.

##### Step 8: ROIs and Masking

One limitation of electroporation is variability in the amount and location of transfection. To minimize the impact of transfection variability on quantification, we compared measurements within three different regions of interest (ROIs): whole limb, the posterior half where wild type ZRS activity is observed, and the zone of polarizing activity (ZPA), the location of active Shh expression found in the posterior third of the limb, within 500μm of the distal tip. To distinguish limb from non-limb tissue, we used the *Roipoly* widget to manually draw a polygon around the limb and export a mask of this ROI. While manual intervention is not optimal, this was the best way we found to distinguish limb from body wall as there is no difference in contrast. The outer edge of the limb was detected with Otsu thresholding. The widget and threshold masks were combined to create a mask that would cover everything in the image except for the limb. To determine the coordinates needed to define the posterior and ZPA regions, we applied *OpenCV*’s contour function to the masked limb image. To locate the extreme left, right, top, and bottom coordinates of the limb, the contour was passed to *NumPy*’s max and min commands. We then obtained the midpoint by subtracting 500μm (349 pixels) from the right extreme point (the distal-most tip of the limb).

Since some limbs are sloped downwards (posteriorly) while others are at a right-angle to the body wall, we determined whether the limb was sloped using an if-else statement, then calculated the average slope of the limb (if it was not zero) by averaging the slope of the top-to-right and left-to-bottom lines. To create the posterior mask, we then used the Path class to create a polygon that would follow the slope of the limb and include only the posterior half. These coordinates were added to a new *NumPy* array of the same shape as the image with the Boolean values inside the polygon being set to True. We then created the posterior mask by combining the posterior polygon mask with the limb mask.

To create the ZPA mask, we first needed to define the lower third of the limb and the point 500μm from the distal tip. To measure 500μm from the distal tip we either used the rightmost point of the limb (right-angled limbs) or the point adjusted for the slope of the limb and subtracted 349 pixels. To define the posterior (lower) third of the limb, we created a line perpendicular to the slope of the limb that transects the midpoint. The coordinates for the perpendicular line are used with the Path class and to make a polygon of the distal portion of the limb. This distal polygon mask is then combined with the limb mask to make the distal mask. We then applied the distal mask to the limb image and used the find_extremes function again to divide the perpendicular line into thirds, again creating a polygon of the lower third of the limb and combining the lower third mask with the distal mask to create the ZPA mask.

##### Step 9: Data Collection

For each method we collected a count of the pixels with a non-zero value, the sum of the values of the pixels and the mean. Additional measurements could be collected, such as finding the geometric center or centroids, but in our application, we found that the area of brightest intensity is impacted by the electroporation technique itself. Such measurements might prove more useful for viral vector systems.

##### Step 10: Putting it All Together

Once users have tested each step of the code and selected which methods to use, this section combines the selected steps to process all images without the additional outputs that are necessary for optimization. Minor edits have been made to streamline the process. This code block creates a *Pandas* dataframe and exports a .csv file with the results. Users can either begin statistical analysis or use the .csv later.

#### 3.1.2 Statistical Analysis

We used the *Pandas, NumPy*, and *SciPy* modules to handle data and visualized data with *Matplotlib* and *Seaborn*. To check for normality, we visually inspected the data using histograms and used the Shapiro-Wilk test for normality from *SciPy*. Outliers were identified using the interquartile range method. After outlier removal, all the data were found to be normally distributed, so we performed a one-way ANOVA (p=2.20×10^-153^) followed by a Tukey’s HSD *post hoc* test from the *Statsmodels* module. A *Jupyter* notebook of the complete statistical analysis can be found on Github at: https://github.com/KateBall/Quantitative_Image_Analysis.

## 4 Results

### Method Comparison: RGB Normalization

We found that 32.5% (±3.8) of the light in GFP images was in the red channel compared to 48.2% (±4.7) in the green channel indicating substantial bleed-through. As a proof of concept, we also decomposed the RFP images to check for GFP bleed-through, we found that 73.5% (±8.8) of the light in RFP images came from the red channel, while only 13.0% (±4.0) was in the green channel, indicating relatively minimal bleed-through. Based on this finding we decided to use only the GFP green channel values for GFP quantification.

Since we split the RGB channels to account for bleed-through, we compared normalizing the following ways:

- RGB/RGB = Intensity of GFP RGB image / Intensity of RFP RGB image
- G/RGB = Intensity of GFP G channel only / Intensity of RFP RGB image
- G/R = Intensity of GFP G channel only / Intensity of RFP R channel only

To accurately represent the value of the experimental reporter (GFP) signal and normalize it to the transfection control (RFP) signal, we compared three methods of data collection and normalization. We compared the intensity values of the RGB images with that of the decomposed green (G) and red (R) -channel only images. We normalized the GFP RGB intensity to the RFP RGB intensity (RGB/RGB), the GFP green channel intensity to RFP RGB intensity (G/RGB), and the GFP G channel intensity to RFP R channel intensity (G/R) (example shown in Figure 3D). In general, there tended to be no significant difference between the RGB/RGB and the G/R method of data normalization (Tukey’s HSD, p≤0.05), but the median value of the G/RGB method was significantly lower than one or the other methods. This seems reasonable since normalizing the GFP G channel intensity to the RFP RGB intensity usually means dividing a small numerator by a larger denominator leading to a lower overall ratio. Based on our RGB channel analysis, since there is a fair amount of red bleed-through into the GFP image, we will use the G/R normalization method in future analyses.

### Method Comparison: Regions of Interest (ROIs)

To account for transfection variability, we collected data within three (nested) ROIs:

- Limb: Includes the whole limb bud, but excludes the body wall and background
- Posterior: The posterior (lower) half of the limb bud, where wild-type ZRS activity is usually detected.
- ZPA: The location of active Sonic Hedgehog transcription, approximately 500μm from the distal (right) tip of the limb bud, and within the posterior (lower) third of the limb.

In general, there are significant differences between groups when used in combination with the Sum method, but not the Mean method (see below) (example shown in Figure 4C, Tukey’s HSD, p≤0.05). The group median increases as the selected ROI decreases. However, there is not always a difference between the limb and posterior ROIs. An argument could be made for using any of these ROIs, though the ZPA is more prone to produce outliers initially. The best ROI method to use could depend upon the scientific question being asked.

**Figure 4:**
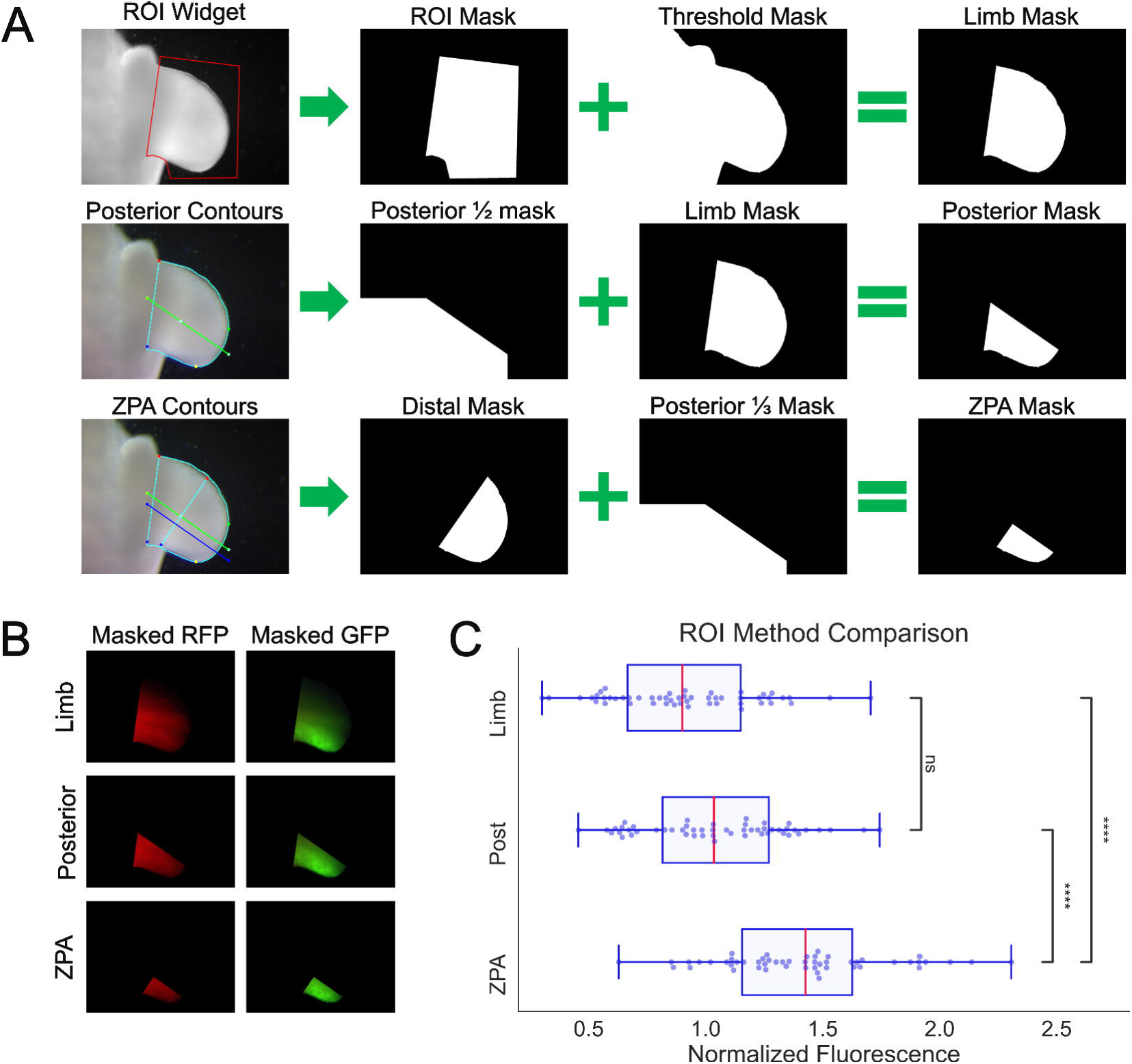
ROIs with hierarchical masking. A) Representative images and masks showing the step to generate the Limb (top row) Posterior (middle row), and ZPA (bottom row) masks, respectively. B) Example of masks applied to RFP and GFP images for data collection (gray scaling and thresholding not shown for simplicity) E) Boxplots representing normalized intensity data collected within each of the ROIs (Tukey’s HSD, ****: p<0.0001, ns: not significant, n=52). Masked RFP and GFP images in A) have been adjusted for lightness (30%) and contrast (10%) using Inkscape 1.2.2.

### Method Comparison: Background Subtraction

To account for variation in background light levels between images/experiments:

- Bg: unaltered.
- Bg removed: each image’s background fluorescence has been subtracted.

Background normalization did not result in a statistically significant difference on normalized fluorescence regardless of the other methods with which it was combined (example shown in Supplementary Figure 1A, Tukey’s HSD, p≤0.05). This may be due to a high level of mechanical consistency and may be microscope- and camera-specific. Although background subtraction did not make a difference in our set-up, we recommend testing this assumption on other set-ups.

### Method Comparison: Sum vs. Mean

The common method in fluorescence image analysis is the use of mean fluorescent intensity (MFI):

- 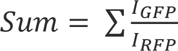
- 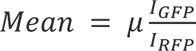

In many cases, taking the mean intensity resulted in significantly higher normalized values than using the sum of intensity (example shown in Supplementary Figure 1B, Tukey’s HSD, p≤0.05). Interestingly, within the ZPA ROI there was no significant difference between using the sum or mean intensity (Tukey’s HSD, p≤0.05). We will likely use the sum of intensity moving forward, as it may reflect greater nuance of spatial information than the mean.

## 5 Discussion

Image analysis has evolved over many decades and a variety of analytic methods have been developed to quantify biological features within images. However, there are few widely used, reproducible, and validated protocols for quantifying features of interest. In this work, we presented a suite of open-source tools in the form of a *Jupyter* notebook to streamline commonly employed image analysis tasks, with the flexibility to compare and optimize various steps. Users can quickly check image saturation, compare noise removal and edge/object detection filters, and create masks using automated and manually-tuned algorithms. Further, we addressed technical challenges such as fluorescence bleed-through and transfection variability, as well as other considerations by comparing four methods of data collection/handling: RGB decomposition and normalization, various ROIs, background removal, and use of intensity sum versus mean.

RGB decomposition showed that on average nearly a third of the light in GFP images came from the red channel. We collected RFP and GFP fluorescence intensity by processing both the original RGB images and decomposed single-channel R (from RFP) and G (from GFP) images. We then compared three normalization methods and found that normalizing RGB to RGB values has a similar outcome to normalizing G to R values. Since GFP images contained a fair amount of bleed-through with the given set-up, using the G/R normalization may best represent reporter activity.

Evaluating fluorescent signal within three biologically relevant ROIs (Limb, Posterior, and ZPA) showed that the smaller the ROI used, the more likely individual data points would be considered outliers. Also, the with decreasing ROI size, the median of the normalized fluorescence increased (Tukey’s HSD, p≤0.05).

Normalizing each image by subtracting its own background did not result in a significant change with this dataset (Tukey’s HSD, p≤0.05), though this step should be considered when using other set-ups.

Comparing sum versus mean intensity showed that often mean intensity was greater than sum intensity, except in the ZPA ROI. This may be because GFP intensity tends to be greatest in the ZPA, and gradually decreases in an anterior direction, while RFP intensity may be relatively bright outside the ZPA. When only the ZPA is considered, there is less variation in GFP intensity. Both measurements of intensity could be used to address different scientific questions.

## Limitations

### Object movement, motion correction

Motion artifacts are common in any imaging application and each investigator deals with this challenge in different ways. Since this approach uses the light image to create the limb mask and then applies it to the RFP image, then subsequently the GFP image, it should be noted that the limb can shift slightly during the manual filter change. Future development could integrate steps from the SIMA package to account for motion artifacts. Also, post-hoc motion artifact correction is possible using tools like Photoshop or GIMP.

### Limb Angle

As limb buds develop, they start outgrowth at a right-angle to the body wall then begin to slope more posterior (downwards). Since the exact timing of development has a gaussian distribution, and sometimes electroporation can cause stunted limbs, we accounted for limbs that are at a right angle and those that are sloped downward. To address this issue, we used the extreme points of the Limb ROI contour to determine whether the limb was at a right-angle or sloped. We then adjusted the definition of the midpoint of the limb using an if-else statement. However, the way the manual ROI is drawn can impact whether the limb meets the right-angled or sloped criteria. To avoid this issue, in drawing the manual ROI, ensure that if the limb is sloped downward, that the left-most point of the limb is not also the top- or bottom-most point.

### File Naming

Currently the code relies on the filename for all identifying information. This is not ideal as it leads to long filenames that are difficult for users to read. We chose this approach because of legacy equipment that was used in the laboratory prior to the development of this work-flow. In the future, the code can be adapted to look for some of this information in the file metadata as well as write metadata out to a file to simplify filenames and maintain metadata for reproducibility. Part of this adaptation is underway as we are developing a class object for specimens that can look for this information either in the filename or in the metadata and store it for the duration of the analysis epoch. This class object will have the attributes: experiment date, embryo number, enhancer, mutant, view, magnification, fluorescence filter, and exposure. Additionally, we can customize the metadata to include other information specific to a given set of experiments.

### Limb size

To segment the limb into the ZPA region, we took images of a ruler on our setup at the appropriate magnification and determined the average number of pixels equivalent to 500 μm (349px) and have used pixel number as the means of measuring 500 μm from the distal-most tip of the limb. However, as mentioned above, limbs can be stunted and sometimes have a width of less than 500 μm which results in an error. In these instances, the ZPA ROI is limited to the width of the limb.

## Future Directions

To make this code useful to a broad audience, we would like to integrate a GUI, likely using Qt or Tkinker, so that coding experience would not be necessary for scientists to benefit from this toolset. The associated code was tested on a limited dataset, however, with a larger test dataset and/or machine learning integration, this could be expanded and adapted. For instance, with ML integration it may be possible to bypass the manual input step, and thus decrease processing time and further limit subjectivity. Machine learning and AI are increasingly becoming central in the biomedical image analysis field (13). This workflow can be used for more than fluorescent reporter assays, but for histopathology, X-ray, computerized tomography, and MRI image analysis. We hope this tool stack is useful to other researchers, and welcome questions and feedback.

## 6 Figure Legends

**Note:** All figures were made using Inkscape 1.2.2 (732a01da63, 2022-12-09), subfigures containing plots, masks, or limbs with contours also used *Matplotlib* 3.5.1, *Seaborn* 0.12.2, and *Statannotations* 0.4.4. Original svg files are available upon request. Any alterations made are indicated in the figure legends and are intended for clarity and aesthetic purposes only. All quantitative data used in this study are collected from raw, unaltered image files.

## 9 Author Contributions

KFB: Writing – original draft, review, & editing, Conceptualization, Data curation, Formal analysis, Investigation, Methodology, Software, Visualization.

JAP: Writing – original draft, Investigation, Methodology, Software, Visualization.

AMC: Writing – review and & editing, Conceptualization, Formal analysis, Investigation.

CUP: Writing – review and & editing, Conceptualization, Investigation.

KCO: Writing – review and & editing, Conceptualization, Funding Acquisition, Resources.

CGW: Writing – review and & editing, Conceptualization, Formal analysis, Methodology, Software.

## 11 Data Availability Statement

The raw data are available at https://github.com/KateBall/ZRS-2025

The original code used in this study is available at https://github.com/KateBall/Quantitative_Image_Analysis under the GNU Public License (GPL, ver. 3).

