## Supplementary Figure 1 for "An Open-Source Image Analysis Method for Quantifying Reporter Fluorescence"

### *Supplementary Material*

### 1 Supplementary Data

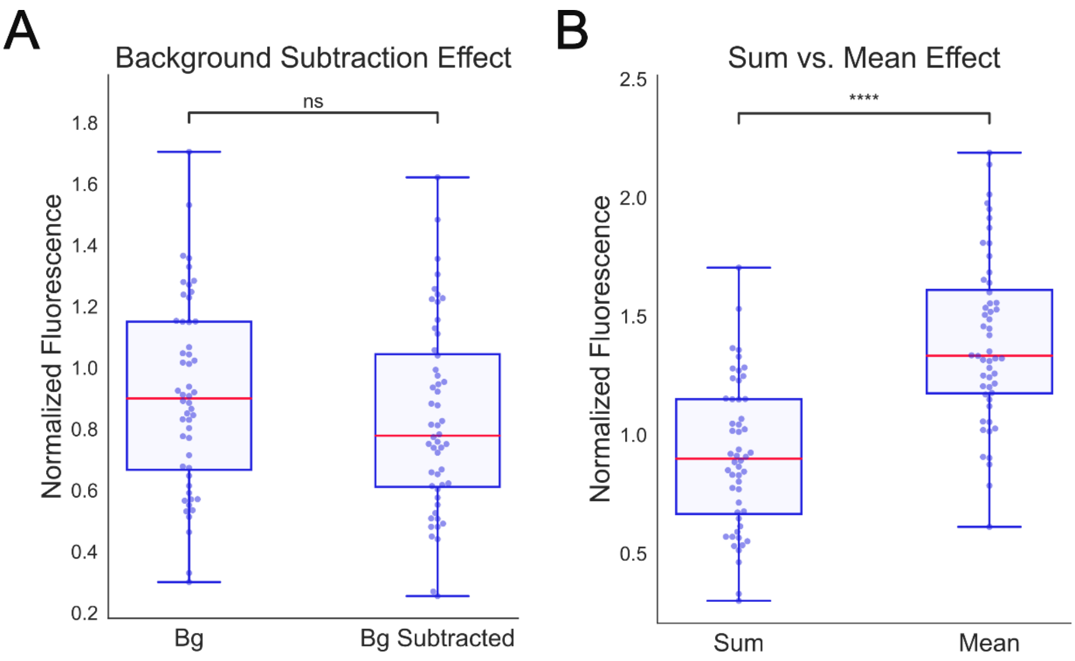


**Supplementary Figure 1: Background Subtraction and Sum vs. Mean.** A) Boxplots representing normalized intensity of either unaltered (Bg) or background-subtracted images (Tukey's HSD, ns: not significant, n=52). B) Boxplots representing either the sum or mean of fluorescence intensity data, GFP normalized to RFP (Tukey's HSD, ****: $p<0.0001$, n=52).
